# Systematic Chemogenetic Library Assembly

**DOI:** 10.1101/2020.03.30.017244

**Authors:** Stephen M. Canham, Yuan Wang, Allen Cornett, Douglas S. Auld, Daniel K. Baeschlin, Maude Patoor, Philip R. Skaanderup, Ayako Honda, Luis Llamas, Greg Wendel, Felipa A. Mapa, Peter Aspesi, Nancy Labbe-Giguere, Gabriel G. Gamber, Daniel S. Palacios, Ansgar Schuffenhauer, Zhan Deng, Florian Nigsch, Mathias Frederiksen, Simon M. Bushell, Deborah Rothman, Rishi K. Jain, Horst Hemmerle, Karin Briner, Jeffery A. Porter, John A. Tallarico, Jeremy L. Jenkins

## Abstract

The assembly of chemogenetic libraries composed of chemical probes provides tremendous value to biomedical research, but requires substantial effort to ensure diversity as well as quality of the contents. We are assembling a chemogenetic library by data mining and crowdsourcing institutional expertise. We are sharing our methodology, lessons learned, and disclosing our current collection of 4186 compounds with their primary annotated gene targets.

## Introduction

Historical screening of compound collections against biological assays along with efforts to characterize the mechanism of action (MoA) of compounds has led to the discovery of many small molecule chemical probes and drugs. This has enabled the construction of smaller focused libraries which, in lieu of throughput have been instrumental toward the identification of potential therapeutic targets, understanding biological signaling, and enabling drug repositioning opportunities.^1,2^ Such libraries can be particularly informative when examining the mechanism underlying phenotypic assays.

The potential of phenotypic screening in disease relevant models facilitates greater potential translatability toward in vivo experiments^3,4^. Retrospective analyses of first-in-class drugs highlight the potential clinical success of drug candidates from systems-based approaches that often originate from phenotypic screening.^5,6,7^ To capitalize on the unique qualities of tool compounds within our own phenotypic screening efforts, we created a strategic workflow to construct a set of probes with the primary goal of broad representation across as many human targets and modalities as possible. Over the course of 6 years, we assembled and grew a dynamic chemogenetic library of chemical probes (a Mechanism-of-Action Box, or MoA Box) using a mixture of data mining and crowdsourcing institutional expertise across a range of cross-functional scientists within Novartis. While a chemogenetic library is not conceptually unique, we wish to share our lessons learned in construction, annotation, and application of our MoA Box so that they may help spur discussions and enable the larger chemical probe community should others wish to undertake parallel efforts. Our efforts have enabled a plated and available chemogenetic library (NIBR MoA Box) for open collaborations with external partners through our Novartis’ facilitated access to screening technologies (FAST) group.

### Informatics driven knowledge automation

The initial NIBR MoA Box construction was guided by general principles^8,9^ outlined for robust tool compounds: 1) demonstrated target engagement; 2) physiochemical properties sufficient for cell-based assays; 3) demonstrated cellular (or organismal) pharmacodynamics effects and 4) appropriate control compounds. A central pillar of our strategy was to be inclusive for the sake of target coverage, with the option to include or exclude probes in a dynamic fashion as internal and external compound–target knowledge evolves. By being inclusive, we acknowledge that hits from our MoA Box screens provide compound–target hypotheses that require additional follow-up validation.

In order to design heuristics to prioritize members, we first needed a large-scale structured bioactivity knowledgebase. HitHub is a robust database and automation system that helps aggregate and integrate chemical and biological data collected from internal, public, and licensed commercial sources, with the goal of supporting informatics-based drug discovery at NIBR (Figure 2). At its most rudimentary level, HitHub facilities informatics applications by aggregating relevant chemical and biological information to a central location. By minimizing the overhead involved in drawing upon information from multiple sources, HitHub facilitates both ad hoc exploration and also provides a robust foundation upon which standard workflows can be developed and shared.

**Figure 1.**
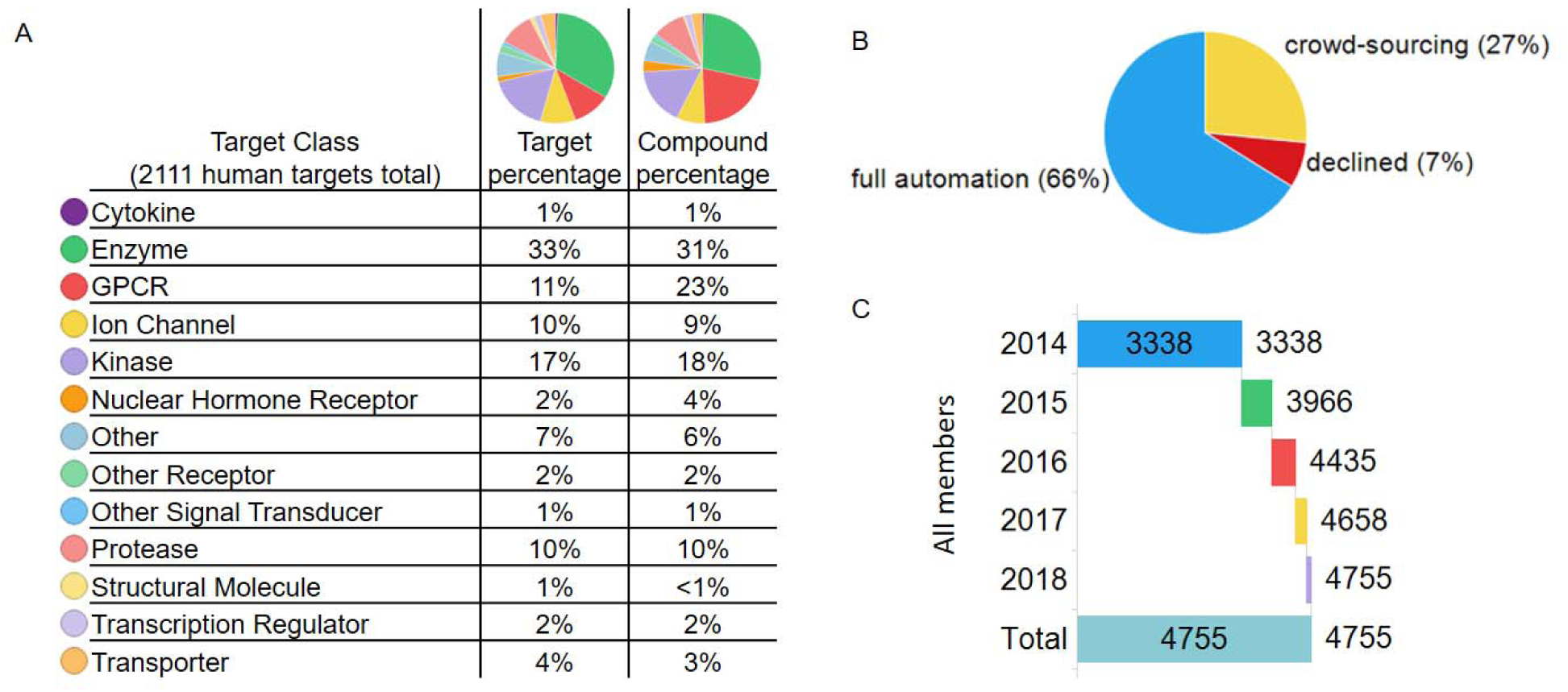
Members in NIBR MOA Box A) Summary of target classes covered as percentage of targets and percentage of compounds. B) Source of compound assembly. C) Growth of members over the past 5 years.

**Figure 2.**
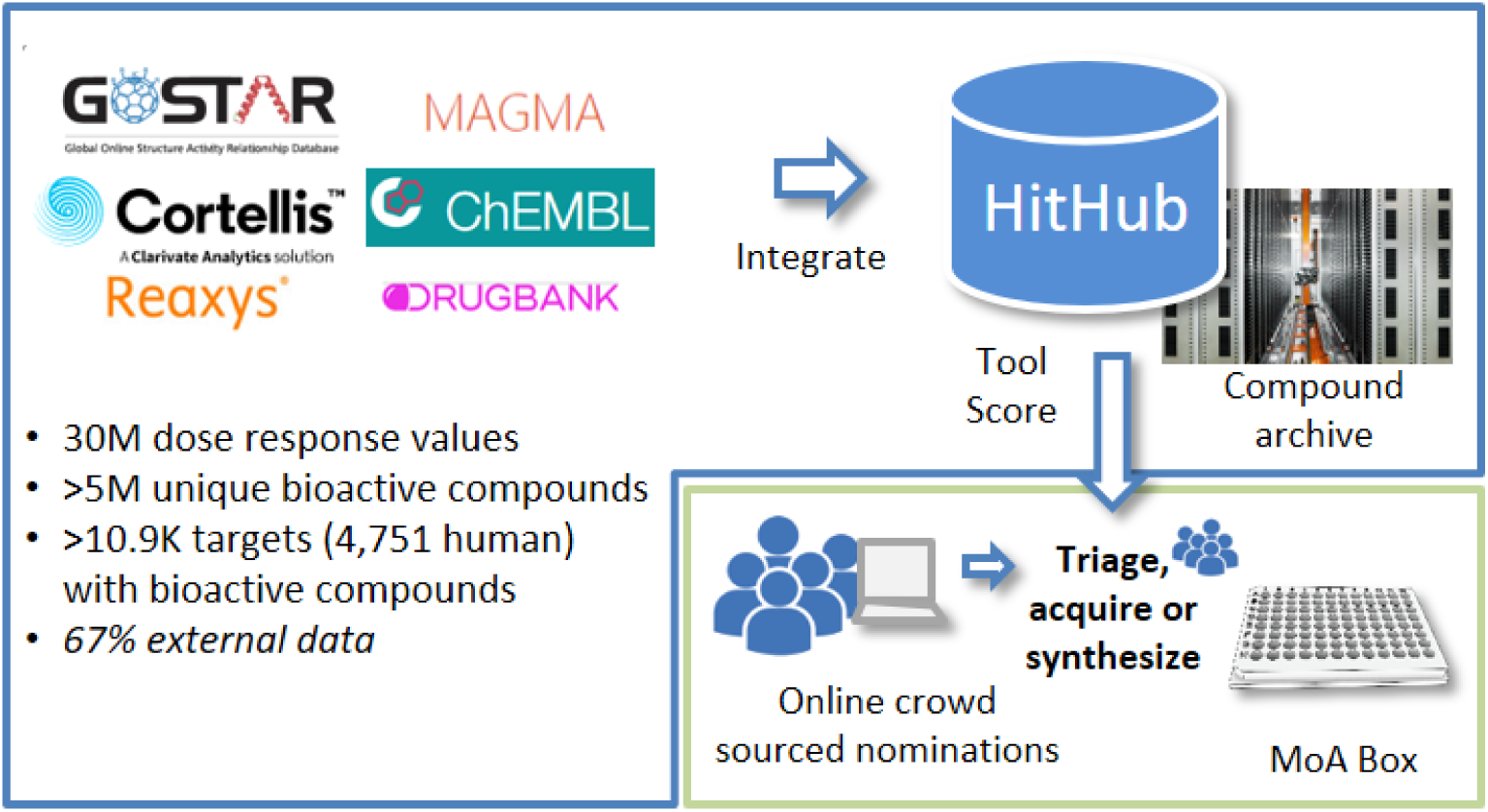
The Novartis HitHub database integrates large-scale compound dose response data in a machine learnable structure. Data sources include public and commercial databases and NIBR data (Magma). Compounds and targets from upstream data sources are normalized to IUPAC InChiKey and NCBI gene IDs and dose response activity values are normalized. In the artificial intelligence workflow (blue), Hithub is queried for a limited number of tool compounds per target with the highest tool scores, with additional rules for chemical properties and availability. In the augmented intelligence workflow (green), scientists make nominations based on experience which are peer-reviewed to determine MOA Box membership.

Often, the most useful of such workflows involve bringing together information from different sources. Such operations, however, require the presence of matching keys shared by the items to be joined. Consequently, an important part of importing data into HitHub involves generating or exposing these keys. To enable joining of data from different sources via common chemical structure, IUPAC InChI Keys are generated for all incoming small molecule data. Similarly, the import process attempts to assign an NCBI gene ID to incoming target data. The presence of these keys increases the integration of the system and multiplies its utility for data analysts. Summary bioactivity tables of all upstream sources are then used to create activity assertions for compound–target pairs.

Having constructed a large-scaled integrated bioactivity data warehouse, we next devised a computational model for prioritization of MoA Box members through a quantitative confidence metric, which we later developed into a Tool Score.^10,11^ In brief, a Tool Score evaluates both the strength of evidence of compound-target modulation and target selectivity. Strength of evidence derives from combining multiple computational inferences or annotations into a strength assertion. Target selectivity parameters were derived by using machine learning to identify descriptors in aggregated compound bioactivity data that maximally distinguish good and poor probes. Using such data-driven quantitative heuristics we can automatically 1) rank potent and selective candidate tool compounds for each individual human target, 2) consider structural diversity of tools, and 3) filter to samples available within the Novartis compound archive. About 2/3 of the current library came from this fully automated workflow (Figure 1B). Where many probes were available per target, we attempted to limit the membership to <10 compounds with a target range of 5, enforcing chemotype diversity using 2D chemical similarity. We prioritized members using Tool Score-like parameters (potency, the amount of papers, patents, or assay data available), pipeline phase reached, and origin, keeping one Novartis-originating compound where available. Compound modality is annotated where available (e.g. antagonist, agonist, destabilizer, etc). In some cases we include probes where the defined mechanism-of-action can be described, but precise resolution to a molecular target among possible isoforms or homologs remains uncertain.

### Leveraging Institutional Insights

The computational approach was extremely powerful for the initial pre-2014 library assembly by large-scale identification and ranking of potential tool compounds, but was limited in the dependence on accessible, warehoused data. Identifying tool compounds from diverse and heterogeneous bioactivity databases was challenging from strictly an informatics computational library design.^12^ Aside from incorrect or unconcise chemical structure assignments, assay target annotations may also be incorrect or misleading (e.g. a protein that is an assay readout or stimulant may be annotated as the assay target). Further the challenge of known and unknown polypharmacology leads to complexity in data analysis (*vide infra*).

We therefore chose to augment our informatics driven approach with the collective institutional knowledge of cross-functional scientists. We believed that scientists with first-hand experience have the best understanding on the most robust and selective chemical probes could enhance and surpass exclusively informatics approaches.^13^ Similar insight-driven crowdsourcing efforts to collect public domain probe molecules have been recently reported.^14,15^ An awareness campaign engaged the Novartis scientific community to cumulate the wisdom of screeners, informaticians, chemists, biologists and pharmacologists and enhance the content of our chemogenetics library. To access the collective insights of the drug discovery scientists at Novartis we created a simple web-based nomination form (Figure 2) open to any individual at Novartis. The nomination page permitted nominations of new compounds (from internal projects or literature), but also to provide expertise and further insights to well-established targets (*e*.*g*. what is the “best” PI3K tool compound) and specific probes (unpublished targets for known compounds). Coupling of our human insight approach aided our informatic strategy by suggesting close analogues of use-restricted probe molecules (*e*.*g*. clinical candidates, controlled substances) or rare compounds (natural products) whose supply was limited. With our insight-driven approach, we could evade stringent parameters criteria for probe inclusion and instead rely on our four simple design principles (*vide supra*). This enabled broader target coverage surpassing the potential liability of *in silico* filtering criteria (activity, selectivity, MW, etc) that would eliminate potentially valuable tool compounds for underrepresented gene targets. Overall, community engagement was vital with 27% of the overall collection originating from the crowdsourced, insight-based approach. As we had hypothesized at the outset of the construction, merging data informatics with human insights yielded a much more robust set of compounds than could have been obtained using either approach independently.

### Continuous cycle of nomination, curation, and acquisition

To maintain perpetual refinement and growth we assembled several dedicated teams to drive nominations, evaluate/prioritize submissions, and ensure acquisition of new chemical probes members. First, we generated a team dedicated to mining the current literature of publications, patents, and conference talks to identify interesting new compound–gene target pairs to incorporate into the NIBR MoA Box. A second triaging team peer-reviewed and evaluated the proposed nominations to help prioritize based on the internal or reported data and gene target(s) coverage in the collection. Probe nominations were declined by the team for failing to meet criteria for competent probes, lack of feasibility of acquiring the compound (rare natural product), or because the supporting data could not be reproduced in house. While the overall number of declined compounds is relatively small in comparison to the total number of nominations (7%; Figure 1B) this activity through a peer-review is vital because to prevent bad tool compounds from being used and misguiding conclusions downstream.^16^ The review team also removed redundancy on well-represented targets (*e*.*g*. PI3K, Bcr-Abl), but enabled the chemotype knowledge to still be captured by taking advantage of a concept that we created so-called “virtual” members which composes 26% of the total library. Such compounds fit all the characteristics of quality probes, but do not need to be included in the library screened in phenotypic assays due to redundancy and can be useful when investigating a target hypothesis.

Physically assembling the NIBR chemogenetic library necessitated an acquisition team to identify internal supply, commercial vendors (∼57% of the set is commercially available), or chemically synthesis the needed chemical probes through partnership with WuXi AppTech, Aurigene Discovery Technologies and Piramal (Figure 1C). On a regular basis the NIBR chemogenetic collection is plated into Echo dispensable plates and used across many exploratory and HTS cellular assays.

## Applications and Data Analysis

Over the course of our efforts, interest in using the NIBR chemogenetics library has sharply increased (Figure 1D). To date the library has been screened in over 300 assays with internal and external academic partners (Box 1) spanning in biological complexity.

Testing the MoA Box collection in these assays has three main purposes: 1) probing novel biology with known MoAs and identifying potential therapeutic targets, 2) evaluating drug discovery flow-chart and counter-assay design 3) repurposing compounds and enriching our knowledge of their bioactivity. To serve these purposes we have applied computational methods to help analyze data and design follow-up experiments. For example, while each chemical probe is included for its primary target annotation, there are often secondary off-targets for each probe, with increased propensity at a higher compound concentrations. These secondary annotated targets and target overlap enable target-phenotype enrichment exercises (e.g. Fisher’s exact test^17^) to help draw out emerging biological trends. Comparing the coupling of biochemical and cellular potency, target-phenotype connection can be further examined (Figure 3). The need for deeper data analysis and the broad use in screening have spurred the internal development of an automated web-based workflow to enable hitlist evaluations and suggest follow up compounds for testing.

**Figure 3.**
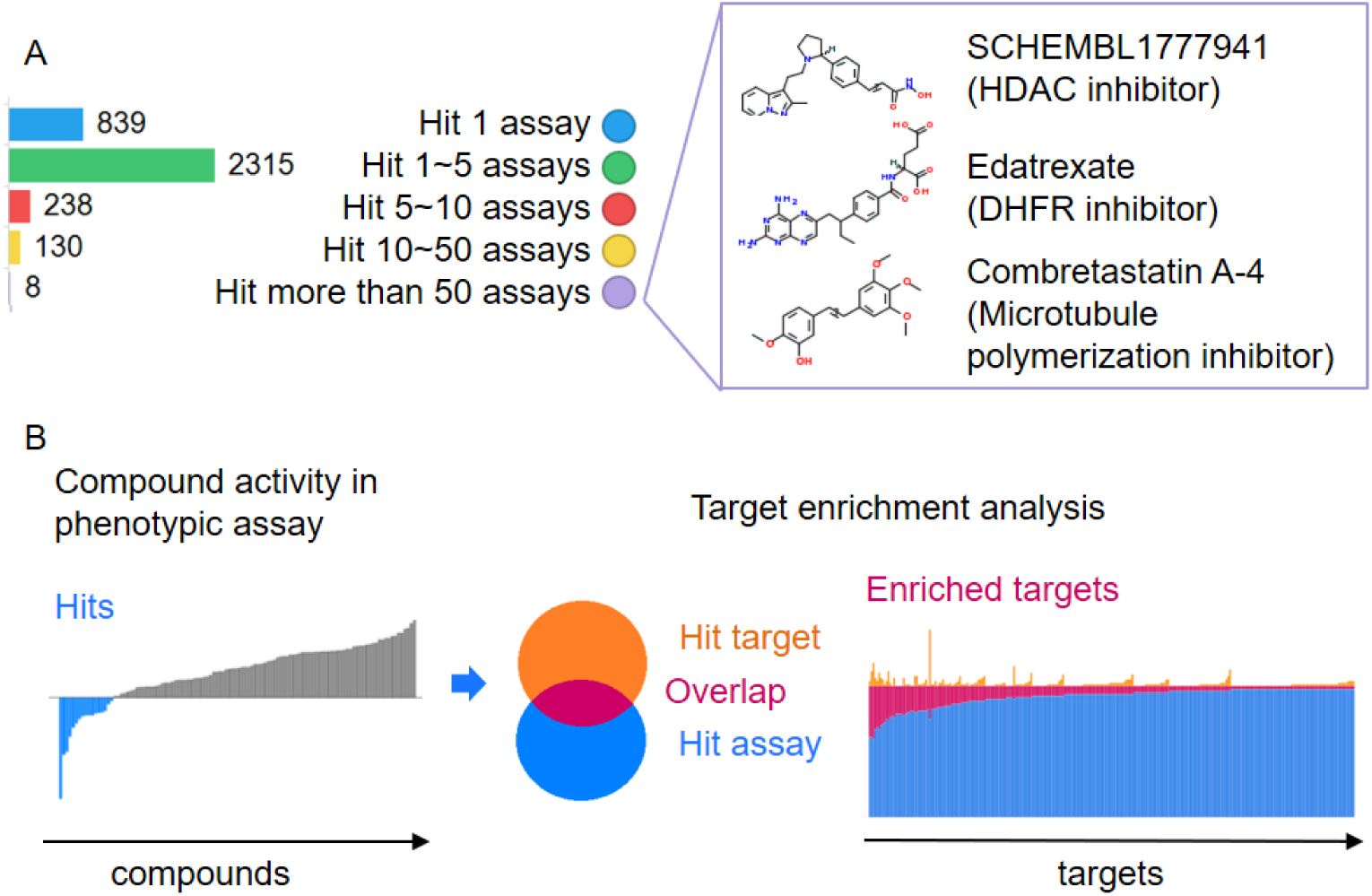
MoA Box screening hits across multiple phenotypic assays A) Number of compounds hitting 1, 5, 10, 50 or more assays. Out of the 8 compounds which hit more than 50 assays, 3 examples are shown. B) Target enrichment analysis. Compound hits in a phenotypic assay are collected and computationally analyzed to identify targets that are enriched among these hits. i.e., if many compounds that are known to biochemically modulate a target are also phenotypically modulating the assay, this target is more likely to be related to the phenotype.

Cross comparison of the MoA box library across a wide range of screens, despite being exclusively comprised of potent cellularly active compounds, has surprisingly not yielded many frequent hitters outside of pan-kinases (e.g. staurosporine) and HDAC inhibitors. On the other end of the spectrum some compounds such as belnacasan, a caspase 1 inhibitor, are hits in very few of the screens conducted to date. This observed screening selectivity highlights the effort to incorporate only high quality chemical probes into the NIBR MoA Box.

To enhance the characterization of the probes and better understand how controlled perturbation of gene targets with small molecules plays out across a diversity of biology the chemogenetics collection is opportunistically profiled with new technologies. For example the library collection has been implemented in a) cell painting assays^18^ to evaluate phenotypic diversity and clustering of phenotypic and target classes b) DRUG-seq, a miniaturization of high-throughput transcriptomic profiling^19^ c) selective cytotoxicity against panels of cancer cell lines to understand predictive drug sensitivity^20^ and d) understanding the mechanisms underlying an assay readout.^21^ Across the many internal pathway screens a few selected examples are highlighted: a) In a Yap-Hippo pathway screen, known targets i.e. ITGAV^22^ were validated and redundant targets, missed by genetic (CRISPR) screening, were identified i.e. TNKS1/2^23^ b) In a Wnt pathway screen, HDAC inhibitors were shown to upregulate the Wnt pathway.^24,25^ c) ALK5 inhibitors were identified as protective of radiation therapy-induced oral mucositis in epithelial spheres.^26^ d) A screen of lung bronchospheres enabled an internal repurposing program to direct basal cell differentiation towards ciliated cells and away from goblet cells.^27^ Overall, we have found the NIBR MoA box collection to be a valuable tool for the typical benefits touted such as uncovering novel biology, focused screening, drug repurposing, etc., but also from a pragmatic aspect of prototyping assays and checking for assay robustness.

### Outlook

Our experience suggests that efforts to expand existing chemogenetic libraries or build new themed collection of small molecules are time and resource intensive commitments. Such efforts need a diverse and committed team to combine informatic data mining and insight driven knowledge.

While we continue to update and improve the NIBR chemogenetics library, we hope that by disclosing the current publicly available portion of our collection (see SI and https://github.com/Novartis/MoaBox) will spur pre-competitive collaboration across academia–Pharma/biotech–commercial suppliers to advance the field of chemogenetic libraries and chemical probes. We have confidence the current disclosed NIBR MoA Box is a solid foundation for interrogating phenotypic screens and is expandable to enable target coverage across the druggable human genome.

#### Box 1.

**Open Collaboration Screening**

Open collaborations at NIBR often involve our facilitated access to screening technologies (FAST) group. External collaborations with the FAST lab scientists is enabled using and in-kind research agreement which provides access to our high-throughput biology instrumentation, informatics expertise as well as a public compound sets that includes the MoA library. This collaboration model implemented in 2016, has since undertaken nearly 50 academic collaborations covering 30 different academic institutes. Many of these collaborations have employed phenotypic assays including the use primary cells or iPS cell types. Highlighted examples of recent collaborations include:

- Discovery that inhibition of p38-MAPK leads to the loss of G1 length and the size uniformity of animal cells.^28^
- Elucidated the role of retinoic nuclear receptor signaling in intestinal organoid morphogenesis.^29^
- Identifed a MAPKAPK5 inhibitor as an antagonist of the STING pathway.^30^
- Unpredictable hit overlaps across human fibroblasts, *Caenorabditis elegans*, and *Drosophila melanogaster* models of the lysosomal storage disorder mucolipidosis type IV (MLIV).^31^
- Demonstrated the development of a highly specific MEK1 stabilization cellular assay.^21^
- Identified agonists of the transcription factor, POU4F3, which induce hair cell generation within the mammalian cochleae.^32^

Through these collaborative partnerships the ability to systematically interrogate complex biological systems has reveal many new biological insights that might be financially or technically limited otherwise. More ready access to such chemogenetic libraries would greatly benefit biomedical research.

## Supporting information

Supplementary Info - LIbrary content and annotation

## Acknowledgements

Ellen Crawford, Tobias Gabriel, Wendi Yajnik, Pierre Nimsgern, Johnny Phan, Xilin Zhou, Catherine Rolando, Estelle Ngo, Nadege Graveleau, Nancy Labbe-Giguere, Amy Calhoun, Laure Bouchez, Paul Selzer, Ludovico Tulli, Christian Parker, Olaf Galuba, Isabelle Adam, Andrea Vaupel, Martin Pfeifer, Christel Guibourdenche, Michael Connolly, Tushar Apsunde, James Neef, Justin Mao, Philip Lam, BoYee Chung, Thomas Caya, Rouwei Mo, Mike Tarselli, Dave Cotter, Mathias Asp, Thomas Veith, Andreas Vallen, Keith Vedananda, Rose Lassos, Micolette Miranda, Sylvain Cottens, Iain Wallace, Ken Yamada, Kayo Yasoshima, David Sandham, Christine Hajdin, Donovan Chin, Vic Myer, Peter Fekkes, John Davies, Adam Hill, Fred Harbinski, Marc Hild, Aaron Wilson, Chris Wilson, Greg Michaud, Markus Schirle, Jason Thomas, Damien MacDougall, Scott Bowes, Caroline Engeloch, Marc Andreae, and all Novartis nominators to the MoA Box.

From Aurigene Discovery Technologies Ltd: Sangamesh Badiger, Prasad Appukuttan, P Venkata Ramana Reddy, Srinivasa Raju, Pritam Biswas, and the Aurigene 020 and 066 teams. From WuXi AppTech Wuhan:Yang Zang, Yanhe Hu, Duan Liu and the WuXi 010 tools team. From Piramal: Tanay Ghoshal and the Neo-12 team.

